# *dropClust2*: An R package for resource efficient analysis of large scale single cell RNA-Seq data

**DOI:** 10.1101/596924

**Authors:** Debajyoti Sinha, Pradyumn Sinha, Ritwik Saha, Sanghamitra Bandyopadhyay, Debarka Sengupta

**Author notes:** these authors contributed equally to this work (Pradyumn Sinha, Ritwik Saha).

## Abstract

DropClust leverages Locality Sensitive Hashing (LSH) to speed up clustering of large scale single cell expression data. It makes ingenious use of structure persevering sampling and modality based principal component selection to rescue minor cell types. Existing implementation of dropClust involves interfacing with multiple programming languages *viz*. R, python and C, hindering seamless installation and portability. Here we present dropClust2, a complete R package that’s not only fast but also minimally resource intensive. DropClust2 features a novel batch effect removal algorithm that allows integrative analysis of single cell RNA-seq (scRNA-seq) datasets.

**Availability and implementation:** dropClust2 is freely available at https://debsinha.shinyapps.io/dropClust/ as an online web service and at https://github.com/debsin/dropClust as an R package.

---

With the advent of single cell transcriptomics, it is now possible to discern phenotypic diversity among seemingly similar cells in complex tissues^1^. Recent development of the Droplet based single cell sequencing technologies has enabled profiling several thousands of cells at a time^2, 3^. Large pool of cells present the opportunity of identifying cell types which are previously unseen due to their limited presence or rarity^4^. Clustering of cells is a primary step involved in single cell expression data analysis. A plethora of techniques exist for this purpose^5^. Majority of these methods struggle with the typical sample size of the data produced by droplet-based technologies^6^. Recently some softwares have been reported for addressing the issue of scalability. SCANPY^7^, bigSCale^8^, DendroSplit^9^ clusterExperiment^10^ and dropClust^6^ are notable among these. The algorithm dropClust was originally designed to attain lower time complexity while not sacrificing on the clustering accuracy. Here we present dropClust2, a memory efficient, monolithic implementation of the original dropClust algorithm with improved execution time and the ability to remove batch effect across datasets, thereby enabling integrative analysis. We hosted the lightweight implementation of dropClust2 on web allowing, for the first time, online analysis of large scRNA-Seq data.

## 1 dropClust and its limitations

Runtime advantage and accuracy of the dropClust algorithm come largely from the use of Locality Sensitive Hashing (LSH) and a de nevo Structure Preserving Sampling (SPS) technique respectively. The existing dropClust software entails various fundamental steps involved in single cell data analysis. These are data normalization, gene selection, clustering, 2D visualisation of cells and differential expression analysis. Moreover, dropClust has been demonstrated to be effective in detecting minor cell sub-populations^6^.

The existing implementation of dropClust suffers from some critical drawbacks, specially with respect to its usability and resource requirements. The existing implementation of dropClust involves interfacing with multiple programming languages *viz*. R, python and C. This hinders seamless installation experience on the user end. Also, dropClust is unfit for standard PCs, due to its memory requirements.

## 2 Improvements

As part of the current release, dropClust has been enhanced in the below areas (refer to Figure S1 for the schematic diagram).

### Selection of Principal Components

In the existing version of dropClust, principal components are re-ordered based on the number of Gaussian mixtures identifiable on the projected data. Bayesian Information Criterion (BIC)^11^ is used to determine the number of modes corresponding to each principal component (Supplementary Figure S4-S6). In practice, Gaussian Mixture Models (GMM) converge slowly. In dropClust2, instead, we use Gaussian Kernel Density Estimator^12^ for convolving a non parametric probability density function (`density` module in R) for projections on each principal component. Probabilities corresponding to 512 random points are sampled using this estimator. Further, a cubic spline curve is fitted (`smooth.spline` module in R) on these 512 points to obtain a continuous approximation of the density function^13^. From the second order derivative of the spline, we record the change in direction of the slope along the curve to determine the number of modes. This approach causes significant speed-up over the previously used GMM technique.

### Introduce Batch effect removal

DropClust2 is equipped with a novel strategy to mitigate batch effects. Our strategy is based on the observation that ranks corresponding to expression values are robust to noise and do not require additional normalisation. However, gene pairs having similar marginal distribution of expression often add noise to the distance computation since their ranks in the individual transcriptomes get flipped highly inconsistently. An algorithm is designed to randomly discard one gene out of such pairs. The algorithm details can be found in the Supplementary Information.

### Disambiguation of post-hoc cluster assignment

DropClust pipeline clusters cells in two steps. First, it clusters a small fraction of carefully sub-sampled (using Structure Preserving Sampling) transcriptomes using hierarchical clustering. The remaining transcriptomes are assigned cluster identities based on the cluster identities of their nearest neighbors. In doing so, previously we followed a simple strategy, wherein the frequent most cluster identity, as represented by the nearest neighbors, was chosen as the identity of a cell that did not participate in clustering^6^. This strategy would forcibly assign cluster identity to a cell even when the highest obtained frequency fails to stand out as a clear majority. To this end, we introduce `conf`, a user parameter (range [0, 1], default 0.5) that applies a cut-off on minimum expected frequency (fraction of the nearest neighbors) of the majority class.

### R package

Existing dropClust uses python module `LSHForest` and C module `Louvain`for nearest neighbor search and community detection respectively. We replaced `LSHForest` with `RcppAnnoy`, which implements a significantly faster version of LSH^14^. C implementation of Louvian was replaced by its R implementation, as part of the popular `igraph` R package.

DropClust pipeline performs dispersion based gene selection at an early stage. This step uses R function`var` through `apply`, which makes multiple copies of the expression matrix, thus incurring significant memory footprint. We implemented a custom function`ColDispersion` in Rcpp that operates on a single copy of the matrix, thus reducing both memory usage and execution time.

### Web-server

We used the Shiny framework to host dropClust2 as a web application accessible through a user friendly wizard, allowing online analysis of large scRNA-seq datasets, as typically obtained from droplet based high-throughput technology platforms (Figure 1 (a), Figure 1(b)).

**Figure 1.**
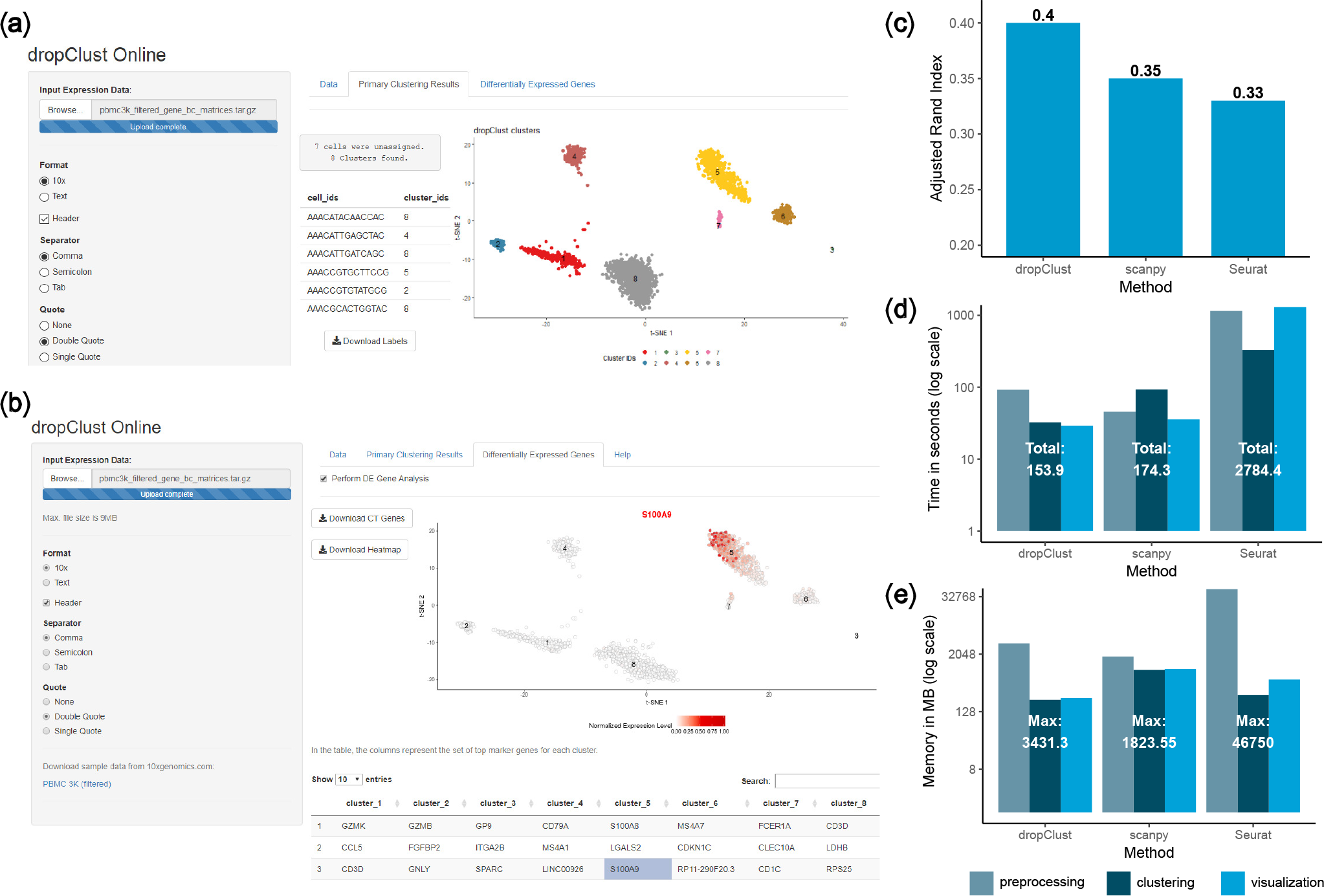
(a) Snapshot of online application after data upload and clustering. The clustering tab contains the clustering scatter plot with predicted clusters, The cluster IDs for respective samples can be downloaded as well. (b) Tab showing the result of differential gene expression analysis. The output contains the clustering scatter plot highlighting the concentration of a selected gene. Genes can be selected from a table accompanying the plot. The table enlists the top 25 genes for each cluster and differential with respect to one-vs-rest basis. (c)-(e) Performance comparison shown on PBMC 68K dataset across methods - ARI, time and peak memory usage respectively.

## 3 Results

Compared to the initial version of the software which took 7 minutes and 20GB of RAM to process a dataset containing 68K single cell transcriptomes of peripheral blood mononuclear cells (PBMCs,^15^), the improved version takes less than 3 minutes and consumes 3.5GB of RAM. The performance enhancement is attributable to the below implementation changes.

1. Use of `RcppAnnoy` R module in lieu of the python version of `LSHForest`
2. Replacement of the `var` module with a custom C function (`ColDispersion`)
3. Substitution of `mclust` (used for GMM) by a spline based peak determination technique

On this data, we recorded an Adjusted Rand Infex (ARI) of 0.4, in contrast to 0.37 obtained using the former implementation. For this we bench-marked the cluster identities against cell type annotations provided by the authors using marker sorted bulk replicates^15^. DropClust2 outperformed Seurat^2^ and SCANPY^7^ by a large margin (Figure 1 (c)-(e)).

Supplementary Figure S3 demonstrates the advantage of batch effect removal on a cohort of ~6K PBMC transcriptomes supplied from two independent experiments.

