## Supplementary Information for "*dropClust2*: An R package for resource efficient analysis of large scale single cell RNA-Seq data"

### Improving *dropClust* scRNAseq analysis pipeline for online deployment

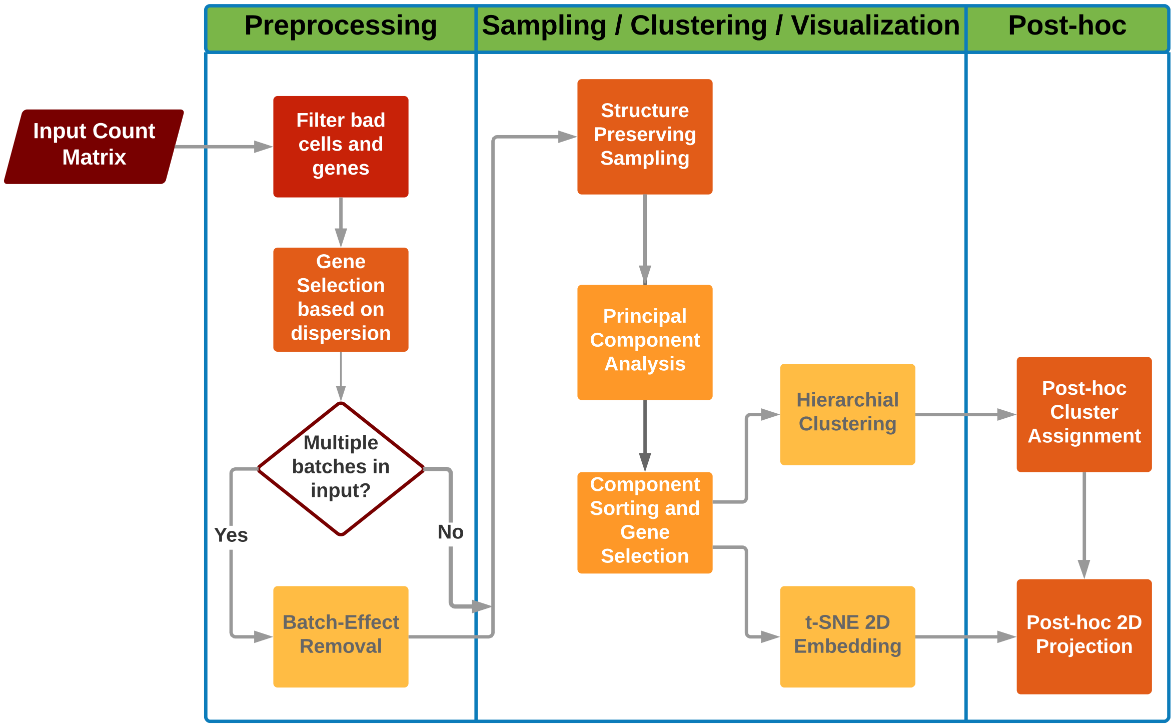


**Figure S1.** Flowchart of dropClust2 pipeline.

#### Batch effect Removal

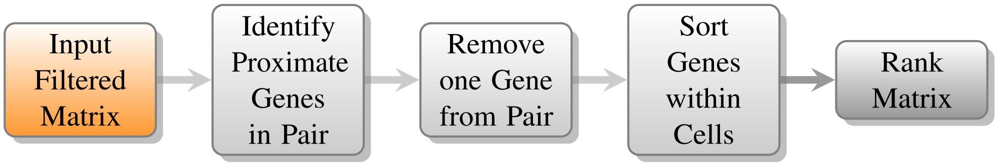


**Figure S2.** An outline of the batch-effect removal stages.

Our approach to minimize batch-effect relies on an assumption that relative ordering of genes with regards to their expressions are preserved across batches. Orderings are also beneficial against changes which affect all genes uniformly, since it is agnostic to magnitudes. However, genes with similar expression values may be attributed distant ranks, which is a unfavourable effect. To combat this we apply a gene removal algorithm, thereby reduce such cases of proximity before conducting the rank based ordering (Figure S2, S5).

##### Gene Removal

The gene removal procedure involves two steps:

**1. Candidate Pair Extraction**: Considering each pair of genes will cause a quadratic complexity with respect to the number of genes. To reduce the number of pairs that are checked, the following method is applied. For each cell, the expression values are sorted and stored in an array. Then the genes belonging to adjacent entries are inserted into the candidate pairs set.

**2. Problematic Gene Pair Identification**: Genes which happen to have close values for a significant number of cell samples are considered to be potentially harmful for the rank based ordering. A pair of genes is identified as 'dangerous' if for more than $k\%$ of the cell samples, the gene expression values of the pair are within $x\%$ of each other. If such a pair is encountered, one of the genes is permanently removed.

##### Rank Based Ordering

The rank based transformation is applied to each cell individually. We consider their gene expression vectors and perform the following steps:

1. Genes are sorted based on their expression values

2. Based upon the resulting indexes, we assign ranks to each of the genes

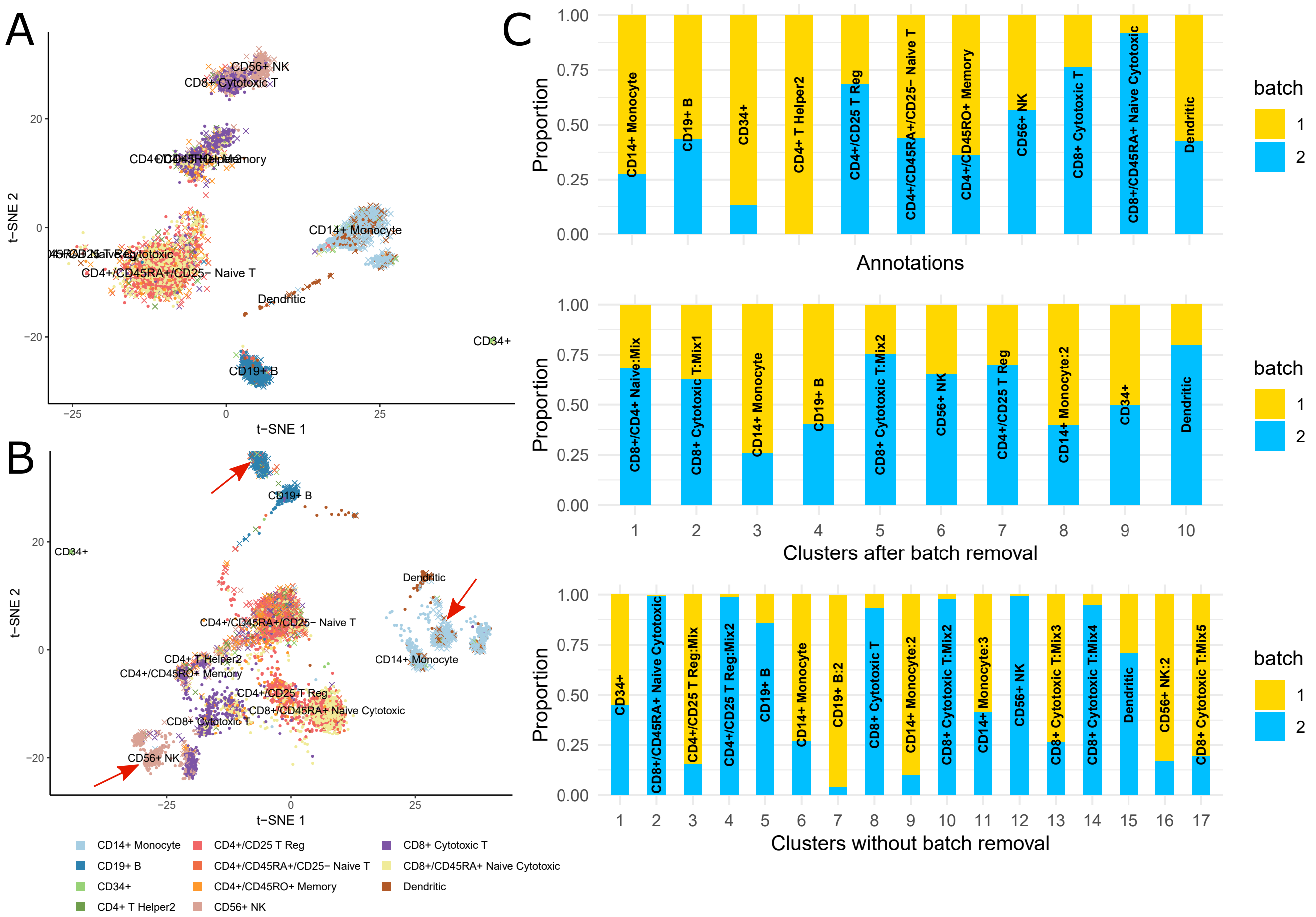


**Figure S3.** Batch effect removal results on a PBMC dataset mixed from two sources. The sources are 1) 3K PBMC dataset from the 10XGenomics.com and 2) 3K PBMC dataset randomly sampled from the original 68K PBMC dataset, also available at the 10XGenomics website. The sources have been referred to as batch 1 and 2 in the figure. (A) Scatter plot, each point represents a cell and coloured by their known annotations. The plot was obtained using dropClust2 after using the batch effect removal strategy. (B) Batch effect removal strategy was not used. The arrows indicate those clusters which originated from almost a single source, specifically B cells, Monocytes and another subpopulation of CD8 Cytotoxic T cells. (C) The barplots indicate the proportion of cells originating from different batches: original annotations, clusters obtained with batch removal and without batch removal. The cluster identifiers were mapped with those cell type annotations indicated by their majority.

#### Principal Component Selection with Results on the PBMC 68K dataset

PCA has been often used for careful selection of genes[1,2]. dropClust has demonstrated that how subjective selection of most informative principal components can improve the gene selection process over traditional variance based component ordering. It observed the components as a multimodal Gaussian mixture, where the modes were contributed by distinct cell types irrespective of their counts. Supplementary Figure S4 illustrates how the first principal component with a large variance masks the presence of the smaller group. The second principal component, on the other hand, exhibits a clear multi-modal pattern and can thereby, easily can distinguish the groups. With this idea, the top principal components ordered by their number of modes was chosen. Supplementary Figure S6 demonstrates how the top components selected by dropClust edges over variance based component ordering. It can be seen that the smaller groups corresponding to cluster identifiers 10-14, is unmasked by the top dropClust components.

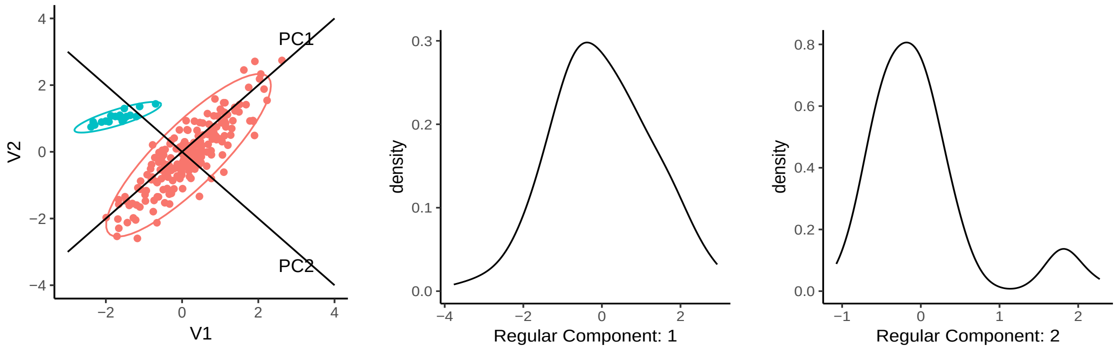


(a) (b) (c)

**Figure S4.** Toy figure to demonstrate variance versus mode criterion. (a) Data containing two groups depicted by their colours, where the larger group dominates the variance in the data. (b) The distribution pattern of the first principal component has a single mode. (c) The distribution pattern of the first principal component has two mode, corresponding to the two groups.

| | **1** | **2** | **3** | **4** | **5** | **6** | **7** | **8** | **9** | **10** | **11** | **12** | **13** | **14** |
|---|---|---|---|---|---|---|---|---|---|---|---|---|---|---|
| CD14+ Monocyte | 21 | 0 | 2 | 1 | 0 | 0 | 1496 | 126 | 792 | 0 | 0 | 1 | 0 | 0 |
| CD19+ B | 11 | 0 | 3708 | 0 | 0 | 0 | 0 | 0 | 0 | 0 | 0 | 81 | 0 | 0 |
| CD34+ | 71 | 2 | 7 | 0 | 0 | 1 | 0 | 0 | 0 | 0 | 164 | 0 | 0 | 0 |
| CD4+/CD25 T Reg | 12541 | 0 | 25 | 1 | 98 | 0 | 1 | 1 | 0 | 213 | 0 | 0 | 0 | 0 |
| CD4+/CD45RA+/CD25- Naive T | 2769 | 0 | 1 | 0 | 1 | 0 | 0 | 0 | 0 | 2 | 0 | 6 | 0 | 0 |
| CD4+/CD45RO+ Memory | 2154 | 0 | 0 | 5 | 223 | 0 | 1 | 0 | 1 | 25 | 0 | 0 | 0 | 0 |
| CD4+ T Helper2 | 7 | 0 | 3 | 0 | 0 | 0 | 0 | 0 | 1 | 0 | 0 | 0 | 0 | 0 |
| CD56+ NK | 6 | 1259 | 2 | 140 | 2 | 2710 | 2 | 0 | 0 | 0 | 0 | 0 | 0 | 0 |
| CD8+/CD45RA+ Naive Cytotoxic | 18999 | 4 | 16 | 167 | 720 | 0 | 0 | 0 | 2 | 10 | 0 | 0 | 0 | 0 |
| CD8+ Cytotoxic T | 617 | 3559 | 44 | 1159 | 872 | 6 | 9 | 1 | 3 | 9 | 1 | 2 | 0 | 0 |
| Dendritic | 188 | 0 | 13 | 0 | 0 | 0 | 333 | 621 | 58 | 0 | 0 | 0 | 79 | 196 |

(a)

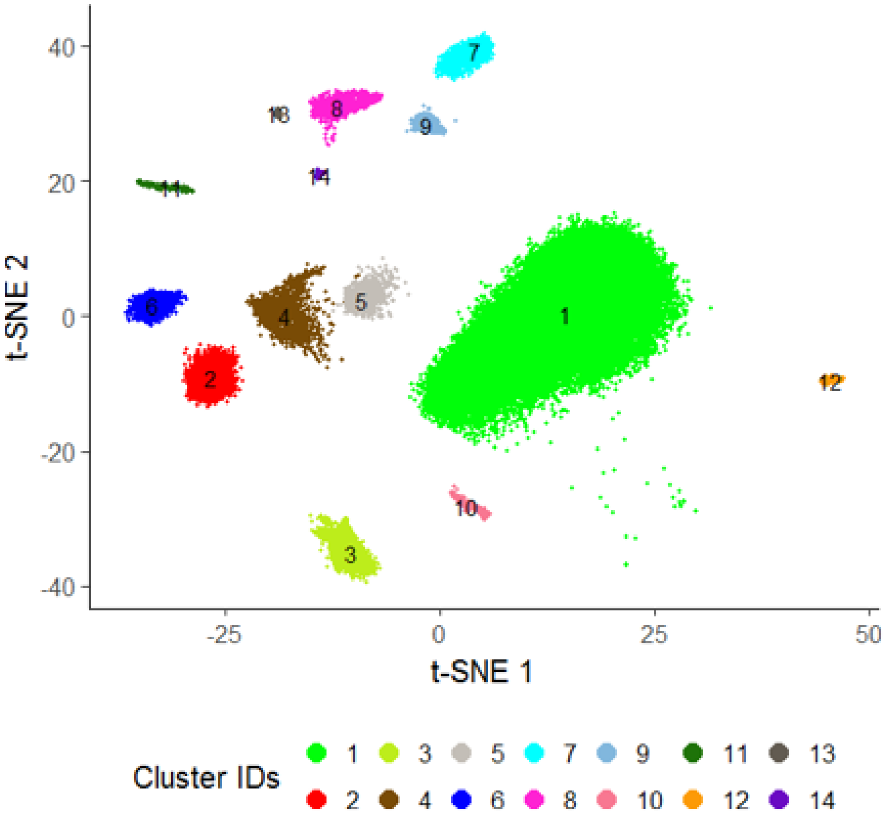


(b)

**Figure S5.** (a) The contingency table depicts known annotations vs dropClust cluster assignments on 68K PBMC data. (b) 2d scatter plot of the clustered 68K PBMC dataset.

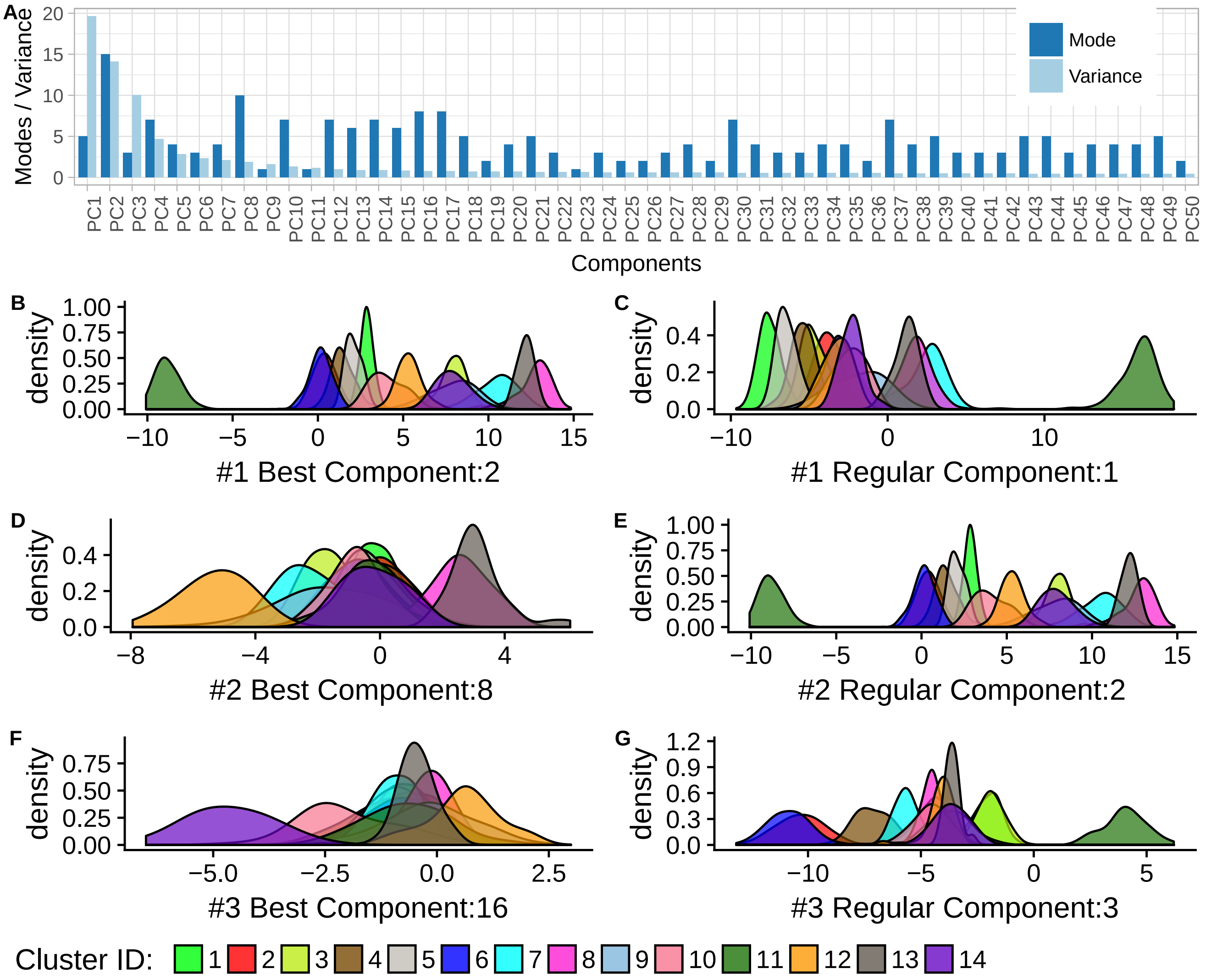


**Figure S6.** Comparing cluster wise distribution of certain Principal components and peaks in overall density curve. (A) The bar-plot represents the number of peaks vs the variance computed for each of the top 50 principal components. (B)-(F) These figures represent the distribution of the predicted clusters in the top three components selected through the dropClust criteria. (C)-(G) The figures represent the distribution of the predicted clusters in the top three components selected through the standard variance criteria. It can be observed that in the former set, more peaks can be distinguished from the first the components. For example the cluster number 10 appears distinctively in figure (F) which is component #16.
